# Methodological Issues with Search in MEDLINE: A Longitudinal Query Analysis

**DOI:** 10.1101/2020.05.22.110403

**Authors:** C. Sean Burns, Tyler Nix, Robert M. Shapiro, Jeffrey T. Huber

## Abstract

This study compares the results of data collected from a longitudinal query analysis of the MEDLINE database hosted on multiple platforms that include PubMed, EBSCOHost, Ovid, ProQuest, and Web of Science in order to identify variations among the search results on the platforms after controlling for search query syntax. We devised twenty-nine sets of search queries comprised of five queries per set to search against the five MEDLINE database platforms. We ran our queries monthly for a year and collected search result count data to observe changes. We found that search results vary considerably depending on MEDLINE platform, both within sets and across time. The variation is due to trends in scholarly publication that include publishing online first versus publishing in journal issues, which leads to metadata differences in the bibliographic record; to differences in the level of specificity among search fields provided by the platforms; to database integrity issues that lead to large fluctuations in monthly search results based on the same query; and to database currency issues that arise due to when each platform updates its MEDLINE file. Specific bibliographic databases, like PubMed and MEDLINE, are used to inform clinical decision-making, create systematic reviews, and construct knowledge bases for clinical decision support systems. Since they serve as essential information retrieval and discovery tools that help identify and collect research data and are used in a broad range of fields and as the basis of multiple research designs, this study should help clinicians, researcher, librarians, informationalists, and others understand how these platforms differ and inform future work in their standardization.

## Introduction

Bibliographic databases are used to identify and collect research data, and therefore function as scientific instruments [1,2]. Studies that rely on these instruments include research on information literacy, bibliometrics/scientometrics, information seeking, systematic reviews, literature reviews, and meta-analyses [3]. These systems, in particular, PubMed and MEDLINE, are also used to inform clinical decision-making in the health professions [4] and construct knowledge bases for clinical decision support systems [5].

Research on search queries that inform the development of bibliographic databases or on how queries influence information retrieval sets were once common lines of inquiry [6] but these have subsided in recent decades [7]. Search query research has largely shifted away from a Boolean model of information retrieval and has focused on ranked-based keyword systems [8] or on database coverage [9–12].

Researchers, librarians, information scientists, and others rely on bibliographic databases to conduct research, to instruct future information and other professionals how to conduct literature searches, and to assist those with information needs to locate and access literature [13–16]. Furthermore, an entire bibliographic universe exists based on bibliographic control that includes standardized rules for description, authority files, controlled vocabularies, and taxonomies to help make searching for information more precise or comprehensive [17,18].

Fine control over bibliographic search and the documentation of search strategies, which are often reported in systematic review research, should allow for the replication and reproduction of searches. In the broader scientific community, the replication and reproduction of research, or lack thereof, has garnered increased attention recently [19,20]. Additional scrutiny has been given to the replication of prior studies [21]. This is true for systematic reviews and other research that relies on citation or bibliographic records, but in this domain, the evaluation of scientific rigor is centered around the reproducibility of search strategies. The Preferred Reporting Items for Systematic Reviews and Meta-Analyses (PRISMA) Guidelines [22]and the Cochrane Handbook for Systematic Reviews of Interventions provide examples of how scholars have recognized the need for the systematic reporting of methods and the organization of review research [23].

Unlike general search engines, bibliographic databases, such as those available on EBSCOhost, ProQuest, Web of Science, Scopus, Ovid, and others rely on structured bibliographic records instead of full text sources to create search indexes. These bibliographic records contain fields we take as meaningful for providing discovery and access, such as author name fields, document title fields, publication title fields, and date of publication fields [17]. In some specialized databases, these records may be supported by controlled terminologies that are database specific or are based on standard knowledge classification efforts, such as the Medical Subject Headings (MeSH) or the Library of Congress Subject Headings (LCSH).

Controlled vocabularies, thesauri, and taxonomies are meant to provide users with a high level of control over the search and retrieval process [24,25]. Some thesauri systems are available across multiple platforms or interfaces [26]. For example, the ERIC thesaurus is freely available at the U.S. Department of Education’s (DOE) ERIC digital library (eric.ed.gov) but also through subscription-based platforms provided by EBSCOhost and ProQuest. Similarly, the MeSH thesaurus is freely available on the U.S. National Library of Medicine’s (NLM) digital library, PubMed, but can be used to search on platforms provided by EBSCOhost, Ovid ProQuest, Web of Science, and others.

Commercial information service companies presumably provide their own access points to bibliographic data from ERIC and PubMed, and the corresponding thesauri, on their own platforms in order to add value above and beyond what the original database providers, like NLM or DOE, have created and provided access to. The added value may be based upon the provider’s unique user interface, its search technologies, its ability to link to library collections via proxy software, its additional database content, or its ability to search against multiple databases on a specific platform in single search sessions.

However, adding value entails some differentiation from the original system [26,27] that may introduce variation in search results. For example, the MEDLINE database accessed through PubMed is defined by the list of publication titles it indexes, the data structure of its bibliographic records, the application of MeSH to the bibliographic records in those records, and the search technologies it implements. When a commercial information service provider also provides access to MEDLINE, it uses that base system, but also differentiates itself from PubMed by providing a different interface, search fields, search operators, and other technologies.

This differentiation among database platforms has long been recognized as important in the systematic review literature in the biomedical sciences, and because of this, the forthcoming PRISMA-S Search Reporting Extension recommends that systematic reviewers report which platform, interface, or vendor is used for each database searched [28]. However, the implications of this differentiation across platforms, with respect to how bibliographic records are queried, are not well understood [29,30]. For example, even though PubMed/MEDLINE, ProQuest/MEDLINE, EBSCOhost/MEDLINE, Ovid/MEDLINE, and Web of Science/MEDLINE are presumably built on the same MEDLINE data file, it is not fully known how the alterations that are made by these vendors impact search and retrieval on their respective platforms. Even when there is some transparency, such as with PubMed [31], these systems are complicated and differences with other systems are not well understood.

Although the choice of database systems impacts potential source coverage and search methods, it is not known how searching the dame database (e.g., MEDLINE) on different platforms might affect source coverage. If searchers used too few databases to conduct literature searches [32–34], then they may miss relevant studies [10,35]. This is especially important in cases where data from past research is collected, analyzed, and synthesized based on published and/or gray literature, such as in systematic reviews or meta-analyses [11,36]. Studies have also highlighted problems in the reporting of search strategies, highlighting incomplete details necessary for others to investigate the quality of a search strategy [37], but this assumes consistency between platforms that provide access to the same database: for example, that using MEDLINE on PubMed is equivalent to using MEDLINE on Ovid, EBSCOhost, ProQuest, or Web of Science. This may have ramifications for those researchers leading clinical trials or conducting bench-side research, and who have to rely on published literature and conduct intensive literature searches when systematic reviews on their topic are not available.

Even if search sessions are methodical and well documented, database systems often operate as black boxes (i.e., the technology is not well documented) and it becomes only possible to infer how different systems operate by comparing multiple implementations [38]. Little is thus known about what actual principles are applied by database vendors in indexing bibliographic records or what specific sets of algorithms are used to rank results when sorted by system-defined relevance. This is commonly known problem among commercial search engines, but it is also problematic in bibliographic databases purchased by libraries [39,40].

Interface, indexing, and retrieval differences also impact reproducibility and replication, which are important aspects of the scientific process, evidence-based medicine, and the creation of systematic reviews [33,41–44]. Although the NLM maintains the MEDLINE records and provides free (federally subsidized) access to them through the PubMed website, they also license these records to database vendors to host on their own platforms. Furthermore, although these systems operate from the same MEDLINE data file, database vendors apply their own indexing technologies and their own search interfaces, and it is possible that these alterations influence different search behaviors and retrieval sets [45,46]. This may be problematic if platform differences are not commonly understood, communicated in vendor reports, or among research team members using them, and if the separate platforms are unable to replicate results based on the same data files that are used across them.

While some studies have included queries that were designed to be reproducible across systems [35], most studies compare queries across systems by evaluating recall and precision on the retrieval sets in these systems [47–49]. However, the focus is not often on the query syntax used even though this has been highlighted as an important problem. One study investigated variations among different interfaces to the Cumulative Index for Nursing and Allied Health Literature (CINAHL) database, and reported reproducible search strategies except for queries that contained subject-keyword terms [29]. In our prior paper (XXXX) we found that queries searched in MEDLINE across different platforms resulted in search result discrepancies after controlling for the search query. This paper extends upon that work with longitudinal data. Here we ask the following research questions:

1. How do search results among MEDLINE-based bibliographic database platforms vary over time after controlling for search query syntax?
2. What explains the variance among search results among MEDLINE-based bibliographic database platforms after controlling for search query syntax?

To answer these questions, our analytical framework is based on the concepts of *methods* and *results reproducibility [50]*. Methods reproducibility is “the ability to implement, as exactly as possible, the experimental and computational procedures, with the same data and tools, to obtain the same results” and results reproducibility is “the production of corroborating results in a new study, having followed the same experimental methods (A New Lexicon for Research Reproducibility section, para. 2). We do not apply the concept of inferential reproducibility in this paper since this pertains to the conclusions that a study makes based on the reproduced methods, and this would largely be applicable if we investigated the relevance of the results based on an information need rather than, as we do, focus solely on the reproducible sets of search queries and the records produced by executing those queries.

## Materials and Methods

We conducted a longitudinal study (October 2018—September 2019) on five MEDLINE-based platforms after two pilot runs in August and September 2018. The five platforms include what is now *legacy* PubMed/MEDLINE (PMML), which is undergoing an interface update that applies new search algorithms [51], ProQuest/MEDLINE (PQML), EBSCOhost/MEDLINE (EHML), Web of Science/MEDLINE (WSML), and Ovid/MEDLINE (OML). The data is based on search result counts for each query in each set and was collected once per month at the mid-point of each month. Twenty-nine sets of search queries were created to search these platforms with each set containing five queries for the respective platforms and for a total of 145 searches per month. The search queries, tested in the pilot studies, were designed to be semantically and logically equivalent to each other on a per set basis. Differences between queries within sets were made only to adhere to the query syntax required for each platform. Table 1 provides an example search set (#010) for October 2018. Each of the queries in this specific set were designed to search for the MeSH term *dementia*, which has two three numbers (MeSH branches), to explode the term, and to limit results by publication date from 1950 to 2015. The last column reports the number of records that were retrieved for each of the platforms for the month and year that data was collected.

**Table 1.** Example set of search queries across the five MEDLINE platforms

|  |  |  |
| --- | --- | --- |
| Platform | Search Set #010 | 10-2018 |
| PMML | "dementia"[MH] AND 1950:2015[DP] | 134217 |
| PQML | MESH.EXACT.EXPLODE( "dementia") AND YR(1950-2015) | 132593 |
| EHML | MH("dementia+") AND YR 1950-2015 | 132599 |
| WSML | MH:exp= ("dementia") AND PY=(1950-2015) | 132590 |
| OML | 1. EXP dementia/ 2. limit 1 to YR=1950-2015 | 132593 |

Our 29 sets of search queries were designed to test the basic functionality of the five platforms in order to compare the results without introducing too much complexity. Therefore, the queries were not designed to test user relevance or user needs, which may range from simple searches to complex, multi-part queries designed for systematic review research. Rather, our queries were designed to test basic functionality with respect to searches that contained some part or combination of the following search fields:

- Keywords
- Specific fields
- MeSH terms with one tree number
- MeSH terms with more than one tree number
- MeSH terms that were exploded

Nine of our search sets included publication date ranges. We added date ranges to test whether search result counts would remain frozen after the end of the publication date and to add an additional control that would help explain differences between the platforms. Most queries also included at least one Boolean operator. All search queries and search result counts are included in the data repository, but Table 2 describes the generalized parameters for each of the 29 sets.

**Table 2.** Generalized search parameters for all 29 sets of search queries

| Search Set Number | Keyword | FieldSpecific | MeSH | Branches | PubDate | Explode | AND | OR | NOT |
| --- | --- | --- | --- | --- | --- | --- | --- | --- | --- |
| 001 | 1 | 0 | 0 | 0 | 0 | 0 | 0 | 0 | 0 |
| 002 | 0 | 0 | 1 | 1 | 0 | 0 | 0 | 0 | 0 |
| 003 | 1 | 0 | 0 | 0 | 1 | 0 | 1 | 0 | 0 |
| 004 | 0 | 0 | 1 | 1 | 1 | 0 | 1 | 0 | 0 |
| 005 | 1 | 0 | 1 | 1 | 1 | 0 | 2 | 0 | 0 |
| 006 | 0 | 0 | 1 | 2 | 1 | 0 | 1 | 0 | 0 |
| 007 | 1 | 0 | 1 | 2 | 1 | 0 | 2 | 0 | 0 |
| 008 | 0 | 0 | 1 | 1 | 1 | 1 | 1 | 0 | 0 |
| 009 | 1 | 0 | 1 | 1 | 1 | 1 | 2 | 0 | 0 |
| 010 | 0 | 0 | 1 | 2 | 1 | 1 | 1 | 0 | 0 |
| 011 | 1 | 0 | 1 | 2 | 1 | 1 | 2 | 0 | 0 |
| 012 | 0 | 0 | 2 | 2 | 0 | 2 | 0 | 0 | 1 |
| 013 | 0 | 1 | 2 | 2 | 0 | 2 | 1 | 0 | 1 |
| 014 | 0 | 2 | 2 | 2 | 0 | 2 | 1 | 0 | 2 |
| 015 | 0 | 1 | 2 | 2 | 0 | 2 | 0 | 0 | 2 |
| 016 | 0 | 1 | 2 | 2 | 0 | 2 | 1 | 0 | 1 |
| 017 | 0 | 1 | 2 | 2 | 0 | 2 | 1 | 0 | 1 |
| 018 | 0 | 1 | 0 | 0 | 0 | 0 | 0 | 0 | 0 |
| 019 | 0 | 2 | 1 | 1 | 0 | 0 | 1 | 1 | 0 |
| 020 | 0 | 4 | 0 | 0 | 0 | 0 | 1 | 2 | 0 |
| 021 | 1 | 2 | 0 | 0 | 0 | 0 | 1 | 1 | 0 |
| 022 | 0 | 0 | 1 | 2 | 0 | 0 | 0 | 0 | 0 |
| 023 | 0 | 0 | 1 | 2 | 0 | 1 | 0 | 0 | 0 |
| 024 | 0 | 0 | 2 | 6 | 0 | 0 | 1 | 0 | 0 |
| 025 | 0 | 0 | 2 | 6 | 0 | 1 | 1 | 0 | 0 |
| 026 | 0 | 0 | 2 | 3 | 0 | 0 | 0 | 0 | 1 |
| 027 | 0 | 0 | 2 | 3 | 0 | 1 | 0 | 0 | 1 |
| 028 | 0 | 0 | 2 | 3 | 0 | 2 | 1 | 0 | 0 |
| 029 | 0 | 0 | 2 | 3 | 0 | 1 | 1 | 0 | 0 |

Note: Column meanings: The *Keyword* column indicates how many keywords were used in the query, not counting field specific keywords, such as document title, journal title, or author name. The latter are counted in the *FieldSpecific* column, which indicates the number of field specific terms used in the query. The *MeSH* column indicates how many MeSH terms were used in the query. The *Branches* column indicates how many trees a MeSH term belongs to. The *PubDate* column is a binary column to indicate whether a query does not include a publication date (0) or includes a publication date (1). The *Explode* column indicates whether a MeSH term was not exploded (0), exploded (1), or in queries with multiple MeSH terms, at least one term was exploded and one was not (2). The *AND*, *OR*, and *NOT* columns indicate a count of how many of these Boolean operators were used in the query. In legacy PubMed, queries require an '*and medline[sb]*' tag in order to limit results to MEDLINE only and to exclude PubMed more broadly. These ANDs were not counted in this column. We did count ANDs when used to join terms or when including publication date ranges in our searches, even for Ovid/MEDLINE, even though Ovid/MEDLINE uses the *limit* operator and not technically the AND operator.

## Results

Our query sets include 100 queries without publication date ranges and nine sets containing 45 queries with publication date ranges from 1950–2015. Thirty-nine of the publication date restricted queries returned different search results from October 2018 to September 2019, indicating potential changes either in the bibliographic records nearly five years after the last publication or potential changes with the search mechanisms used among the MEDLINE platforms. This discrepancy yielded insights for the differences we found across the search sets, and in the following sections, we describe some major themes of these differences among these platforms.

### Macro and Micro Views of the Data Reveal Different Trends

A macro examination of the search sets restricted by publication date (search sets #003–#011) mainly indicated that there were only substantial differences in total search result counts over time between platforms within each set, for example, between WSML and PMML (Figure 1, top left plot). This macro view would appear to indicate that although there are differences among platforms, the comparable tends are reliably consistent across time on a per query, per platform basis. However, a re-scaled, side-by-side comparison of result counts per platform indicates more variation within the platforms themselves (Figures 2–4). This illustrates that platforms are not internally reliable across time on a per query basis. Furthermore, there is also no apparent difference in the growth trends between queries that are restricted by publication date and those that are not restricted by publication date (Figure 5). Figure 5 illustrates the annual growth of records over the year for sets with queries restricted by publication dates and sets of queries with no publication date restrictions and shows that queries restricted by publication dates continue to return new results years past the end of the publication date range.

**Fig. 1.**
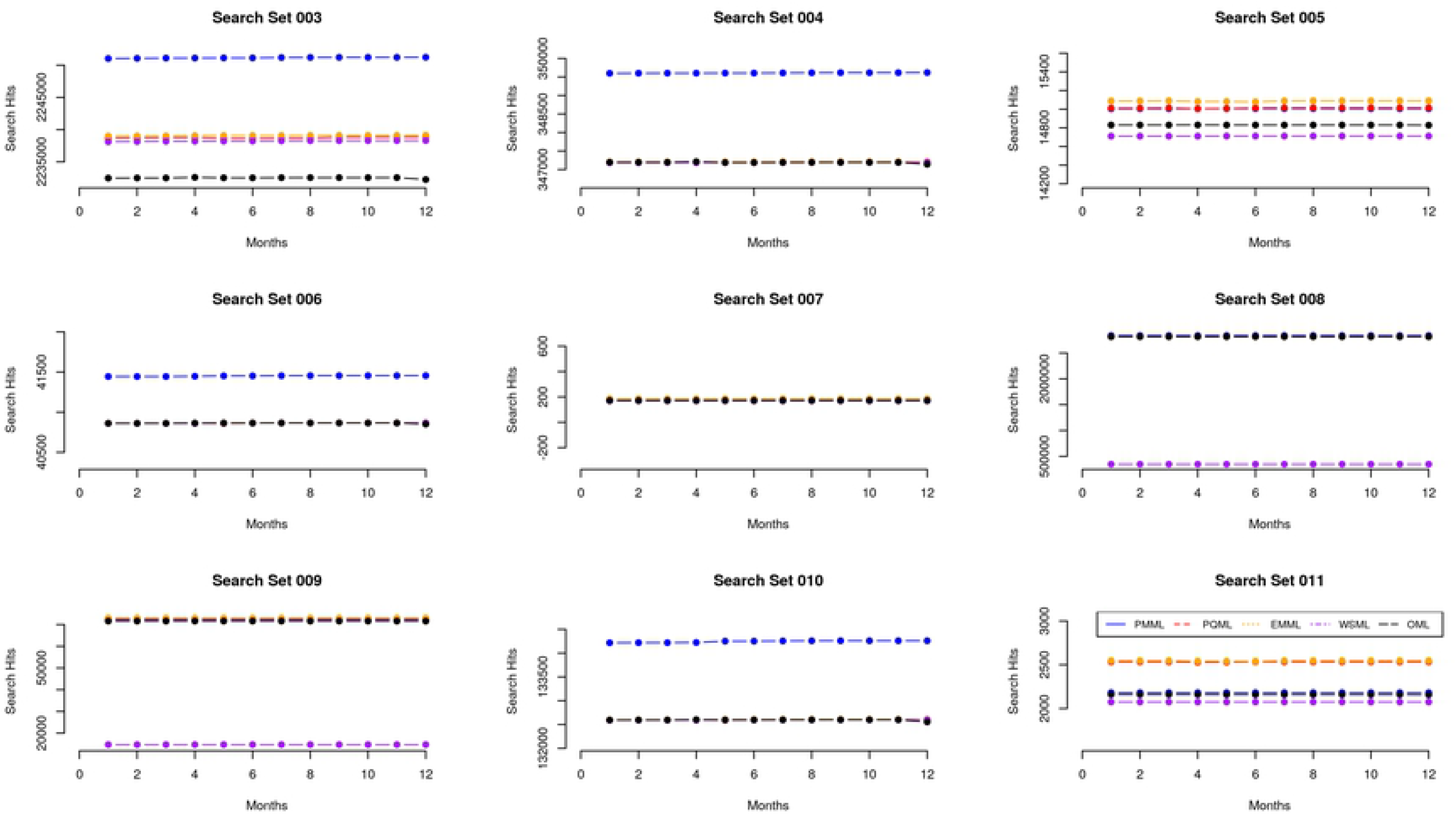
A macro scaled view of search result counts for eleven search sets restricted by publication date. Each plot indicates a separate search set.

**Fig. 2–4.**
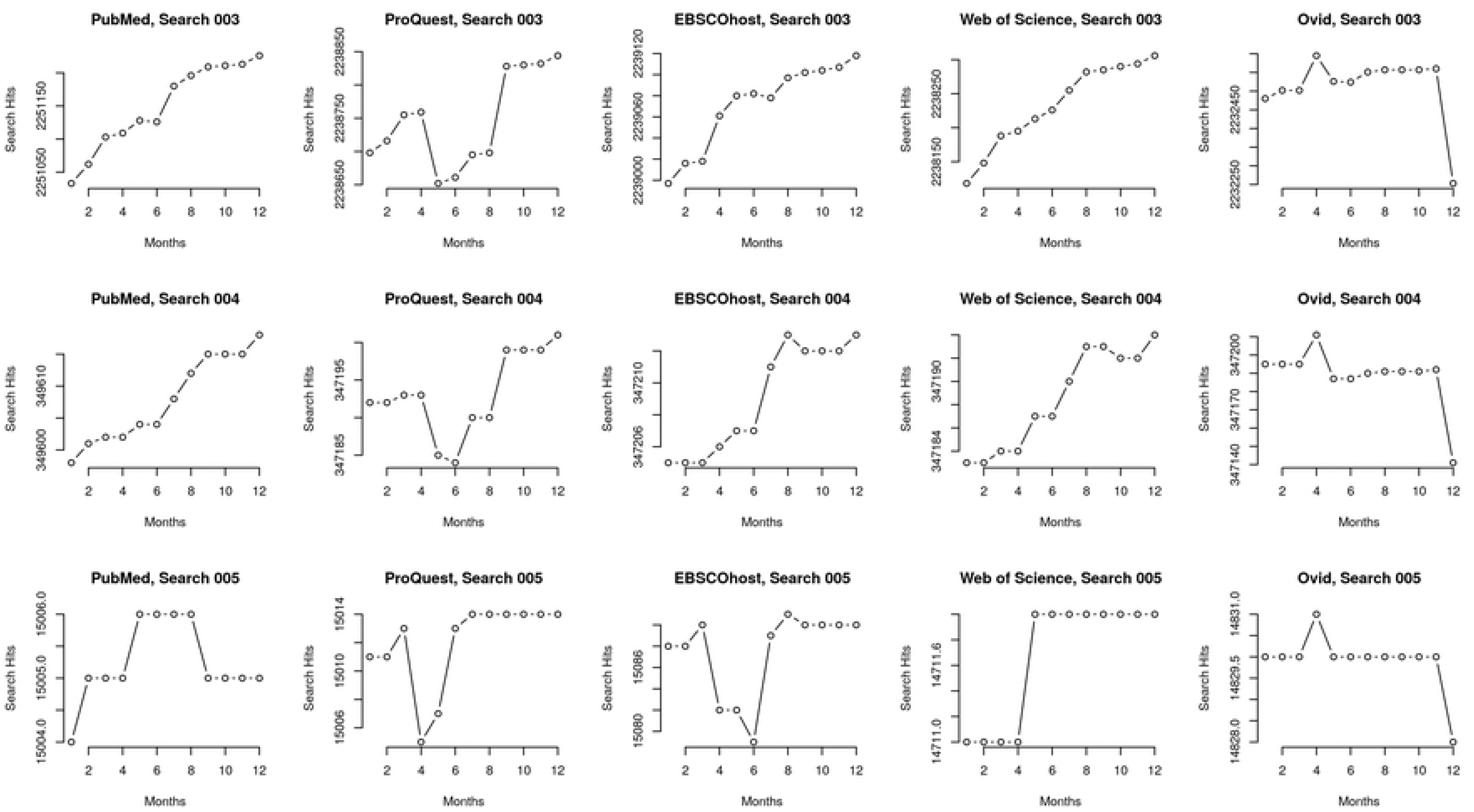
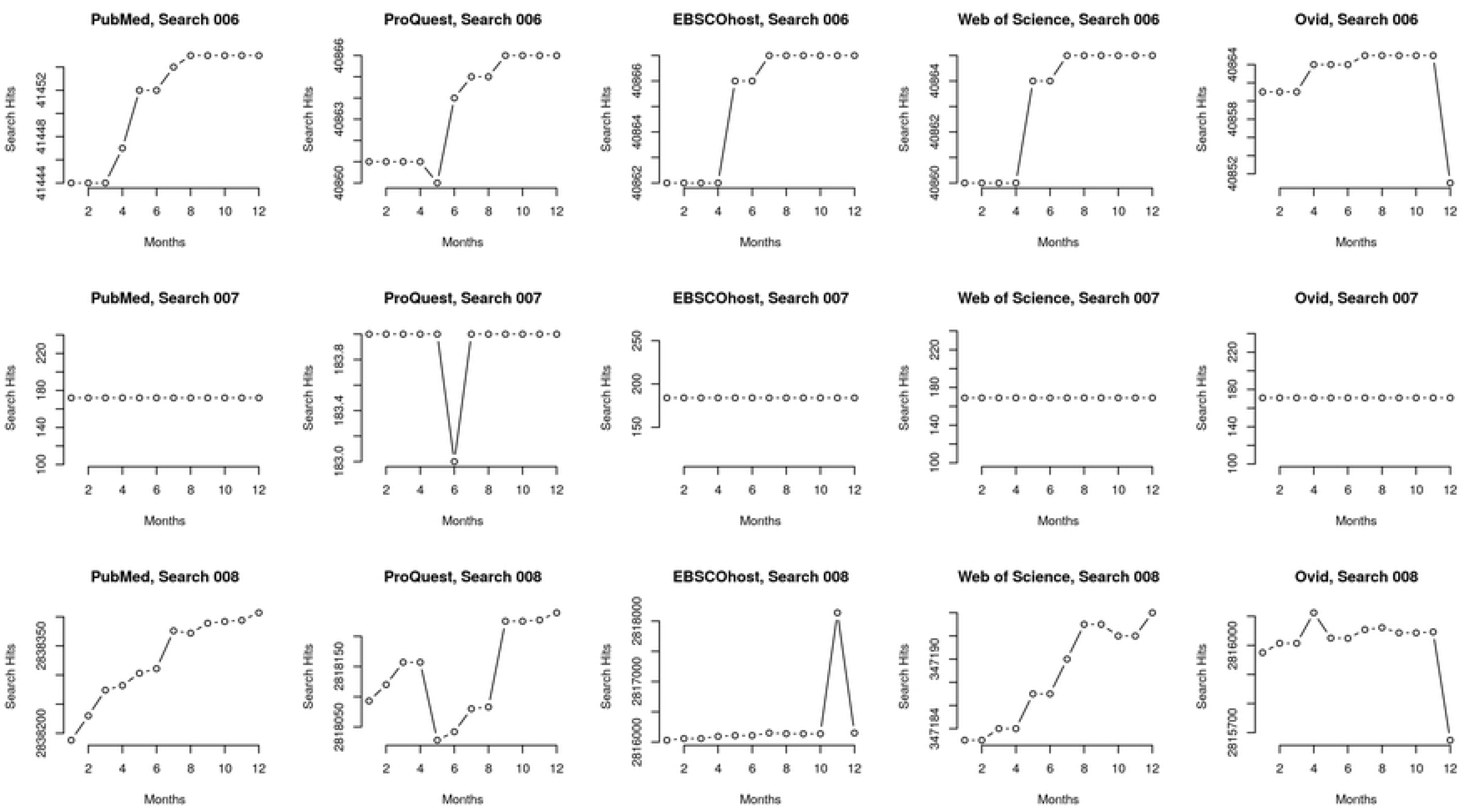
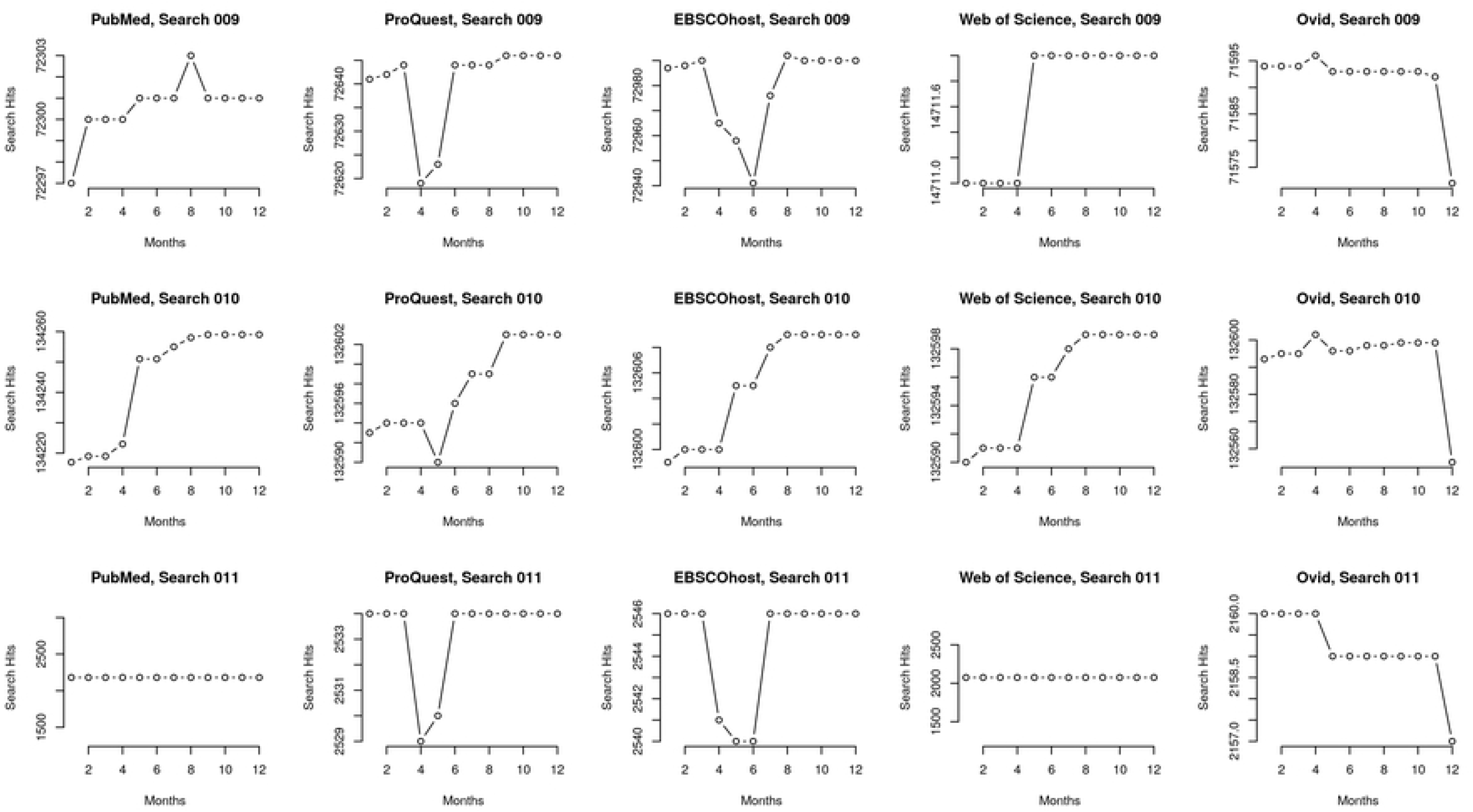
A micro scaled-view of search result counts for eleven search sets restricted by publication date. Each row indicates a search set.

**Fig. 5.**
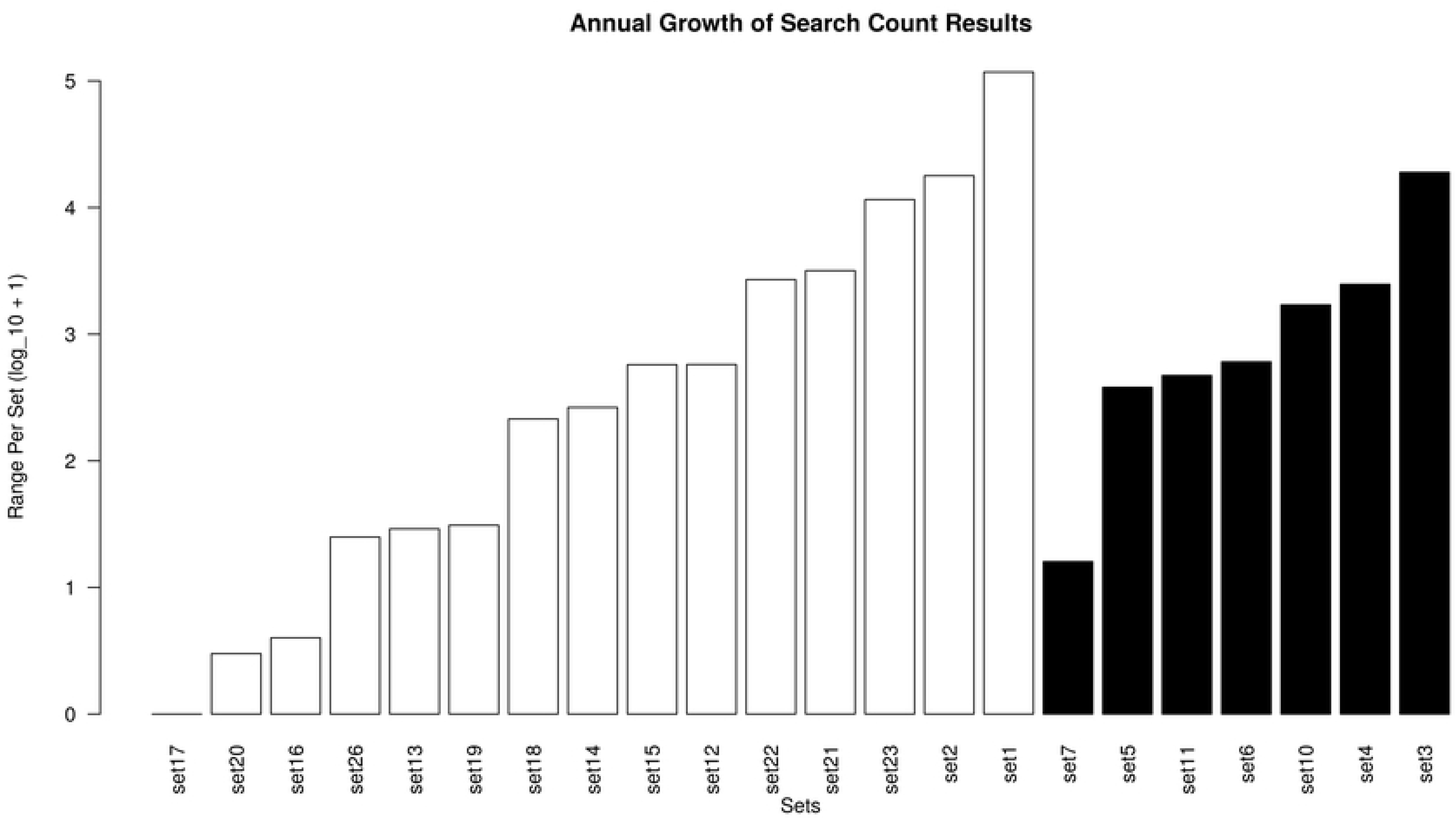
A side-by-side comparison of the growth of search result counts for search sets that were not limited by publication date (white bars, on the left) and search result counts for search sets that were limited by publication date (black bars, on the right).

### Online-first and Print Publications Reduce Effectiveness of Bibliographic Control

To help identify an explanation for the changes in search results for queries that were restricted by publication dates, we compared two query sets, query #002 and query #004, which were both designed to search for a single MeSH term (“neoplasms”), non-exploding, but differed in that query #004 is publication date restricted. Hypothetically, searches for MeSH terms should not be impacted by changes in search mechanisms since the search process for a controlled term is based on whether a record contains the term or not. The grand median change over the year in search result counts for query #002 among all five platforms was 16,551 records (max: 17102; min: 15933), indicating the hypothetical annual growth in literature attached to this term since this query was not restricted by publication date. The grand median change in search results for query #004 among all five platforms was 17 records (max: 70; min: 8) for query #004. Since this query set was restricted by publication date, this indicates hypothetical changes that are not related to literature growth but to changes in either the database search indexes or the bibliographic records, four years after publication.

Furthermore, since all platforms reported different search result numbers in this query set, this indicates that the five platforms are indexing different versions of the same MEDLINE file, or that the platforms index the basic MEDLINE file differently based not on the MeSH term but on the publication date field. To test this, we traced a record from the #004 query results. The record we investigated was chosen because it was part of the retrieval set for a query that was limited by the publication date to 2015 but which the record indicated it was published in 2019.

According to PubMed documentation, the default publication date field [DP] or [PDAT] includes the date of publication for either the electronic or print version [52]. An investigation of the chosen record from the search results for #004 in the PMML set [53] shows a bibliographic record with a long publication history. The PubMed record indicates that the record was added to PubMed in 2015 but not entered into MEDLINE proper until 2019 (See: https://www.ncbi.nlm.nih.gov/pubmed/26700484?report=medline&format=text). On the journal (BMJ) web page, there are two versions of the article—an “Online First” version for the article that was issued in 2015 (See: https://spcare.bmj.com/content/early/2015/12/23/bmjspcare-2014-000835.info?versioned=true) and an “online issue” version of the article that was issued in 2019 (See: https://spcare.bmj.com/content/9/1/67.info). The journal article’s publication history on its web page states that the 2015 version of the article is the online first article, and the 2019 version is the publication date for when the article was assigned to a volume and issue and when it then appeared in print and was added to MEDLINE based on the bibliographic data attached to the 2019 version. On the journal’s site, there are thus two versions of this article. On PubMed, there is one record for this article with two publication dates because of versioning.

The above record indicates problems with bibliographic control and dependency on journals to maintain bibliographic records that are complicated by two sequences of publications: online publications that precede print publications for the same article, or Online First publications that preceded publications that are attached to volume and issue numbers. The latter are further complicated by the versioning of articles based on publication history and that include versions prior to their official publication dates when they are assigned to volume and issue numbers and then added to MEDLINE. This problem with bibliographic control impacts search results across the platforms. The BMJ article described above does not appear in the search results among the other four MEDLINE platforms for query set #004. This confirms that the PMML platform captures the electronic publication date by default even though the record was not entered into MEDLINE proper until the print publication date which, in this case, was four years after the electronic publication date and even though the query was restricted to MEDLINE. Neither PQML, EHML, WSML, nor OML behave in this way. PQML and WSML only offer the ability to search by a single publication date limit, which seems to be defined by the e-publication date, and these platforms do not offer the ability to search by other date fields. EHML and OML offer more control over a variety of date fields but apparently the default publication date field is not inclusive of print publication dates between these two platforms, like it is in PMML.

### Reproducible With Specific Field Searches

We found that the sets of queries that returned nearly equivalent search result counts were queries that included additional specificity, regardless if the queries were restricted by publication date. Query #013 included two MeSH terms, the first one non-exploded and the second one exploded, that were connected by one Boolean NOT, plus one document title term. All five platforms returned results that were within a range of 3 records, taking into account the range of results per platform and then among platforms over the annual period of data collection. This relative consistency across platforms was found in other search sets that included additional, specific field searches. For example, query set #018 performed a single author search for an author who was chosen because they had published in journals indexed by MEDLINE during the middle parts of the 20th century. The range of records that were returned varied over the months and numbered within a range of 15 records among the others. However, when a MeSH term was added to the author name search (search set #017), chosen because the author had published papers that had been indexed with the specific MeSH term (“neoplasms”), all five platforms returned the same count of records for all twelve months of data collection.

### Database Integrity Fails

Aside from issues with bibliographic control due to online versioning, and with differences in indexing, several of the platforms returned results that appear as outliers compared to the others within a set. The query in search set #008 included one MeSH term, on a single branch, exploded, with a publication date restriction. PubMed/MEDLINE returned a range of 219 additional records across the months. ProQuest/MEDLINE returned a range of 211 records, and Ovid/MEDLINE returned a range of 438 records. However, EBSCOhost/MEDLINE returned a range of 2108 records, and although Web of Science/MEDLINE returned a range of only 11 records for the time period, it also returned an average of 2491138 fewer records than PubMed/MEDLINE. We could find no discernible reason for this discrepancy. Search set #010 and search set #023 both included a single MeSH term, two branches, exploded, and although search result counts were different among these platforms within these sets, the differences were not as extreme, perhaps then indicating a problem with how Web of Science/MEDLINE explodes single branch MeSH terms.

There were two query sets where one platform failed to return any results. In sets #024 and #028, Web of Science/MEDLINE returned 0 results across all twelve months, even though the syntax of the queries were correct and one of the queries had returned results in a pilot test but then dropped them in subsequent tests. Additionally, in sets #025, #027, and #029, the Web of Science/MEDLINE search result counts were initially within a reasonable range of the other platforms in the respective sets, but then diverged substantially. For example, in search set #025, Web of Science/MEDLINE returned a maximum of 13021 records and a minimum of 12652 records from October to April. However, the same query returned a maximum of 629 records and a minimum of 619 records from May to September, indicating a drop of over 12000 records for the same search query. In search set #027, Web of Science/MEDLINE returned search counts that were different but comparable to the other four databases, but then in May again, the counts increased by nearly 40000 records and remained within that range until the end of data collection. For search set #029, the search result counts were again within range of the other four databases through April, but then in May and until the end of data collection, the query no longer retrieved any records. The only pattern among these three queries was that the sudden changes in search result counts occurred in May.

### Differences in Database Currency

The time it takes to import the overall PubMed file among platforms also impacts the retrieval sets. On January 24, 2020, the US National Library of Medicine recommended an interim search strategy for searching PubMed for literature related to the Covid-19 virus [54]. Their recommended search query searched all fields for the term *2019-nCoV* or for the terms *wuhan* and *coronavirus* in the title and abstract fields:

#### 2019-nCoV[All Fields] OR (wuhan[tiab] AND coronavirus[tiab])

We modified the search strategy to use it among all platforms and queried the PubMed platforms at irregular intervals. Results show that for the PubMed data file, generally, all of the platforms return different results for the same query for new literature. Results also include three versions of NLM’s interface to PubMed. Two versions are for legacy PubMed but show result counts for when records are sorted by Best Match or Most Recent, since legacy PubMed applied two different sorting algorithms in this version of PubMed [31]. We also show search results for the new version of PubMed, which does not apply different sorting algorithms for Best Match or Most Recent, but which does report different search counts than both legacy PubMed results. As Figure 6 illustrates, these different platforms for PubMed retrieve newly added records at different rates, likely because they receive and incorporate the PubMed data file at different times (Figure 6).

**Fig. 6.**
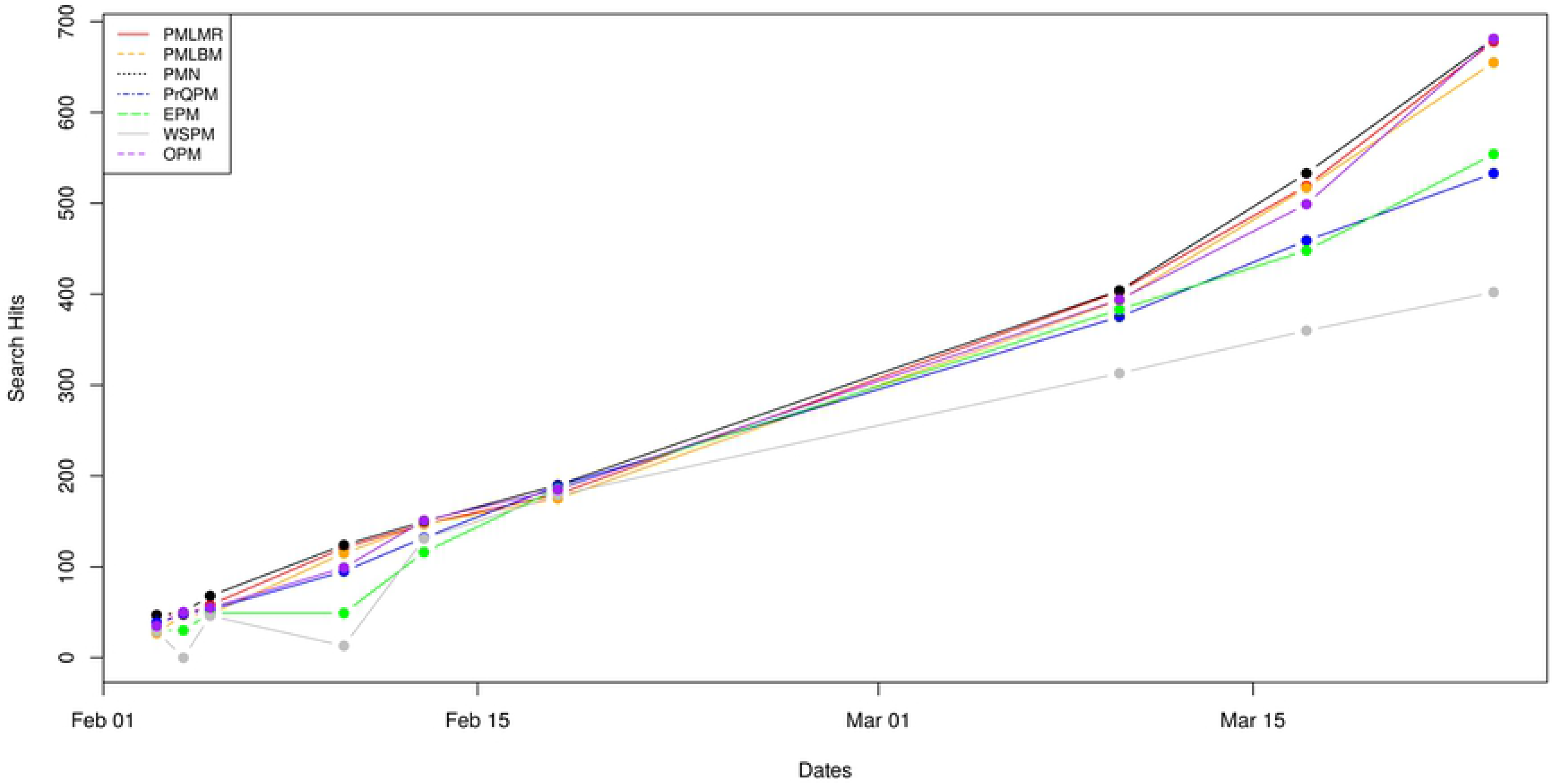
Search result count differences for COVID-19 related searches across PubMed based platforms. PubMed Legacy Most Recent Sort (PMLMR), PubMed Legacy Best Match Sort (PMLBM), PubMed New (PMN), ProQuest PubMed (PrQPM), EBSCOhost PubMed (EPM), Web of Science PubMed (WSPM), Ovid PubMed (OPM)

## Discussion

MEDLINE is the National Library of Medicine’s premier bibliographic database that contains more than 25 million references to journal articles in life sciences with a concentration on biomedicine. The subject scope of MEDLINE is biomedicine and health, broadly defined to encompass those areas of the life sciences, behavioral sciences, chemical sciences, and bioengineering needed by health professionals and others engaged in basic research and clinical care, public health, health policy development, or related educational activities. MEDLINE also covers life sciences vital to biomedical practitioners, researchers, and educators, including aspects of biology, environmental science, marine biology, plant and animal science as well as biophysics and chemistry [55].

### Methods and Results Reproducibility

Overall, we found several issues that impact the unevenness of search results across these platforms and therefore their use as reproducible scientific instruments. Due to differences in search fields across MEDLINE platforms, such as with the publication date field, in developments in publishing, such as online first versions of articles versus volume and issues number versions, in the ability of databases to behave consistently over time, and to differences in updates of the source file across platforms, it is difficult to construct queries that perform consistently alike across systems and to get results that are consistent across systems.

Specifically, we found that queries restricted by publication dates continue to return new records years past the limit on the publication date range. Data from this study begin to provide some explanation for this variance. First, the growth of “online first” publications seems to have complicated the traditional bibliographic record for journal articles which, in part, relies on the relationship between an individual article and its volume and issue. The inclusion or absence of metadata elements such as these are perhaps resulting in the creation of multiple records. In some cases, changes to the original record has meant changes to the MeSH indexing. Additionally, although all platforms provide a search field or a way to limit search results by publication date, not all do so at the same level of detail. While simple publication date searching may have been sufficient in a print only age when there was only one publication date, it is not sufficient in an age when articles are published multiple times, via electronic publication dates and via print dates with volumes and issues. The implication of each of these examples is that records risk being dropped or added to time-restricted searches. Studies that rely on replicable search strategies are at risk of being inherently flawed.

We further found that queries were more likely to return comparable search result counts when they included multiple and specific field terms, such as queries combining keywords appearing in a journal title or an article title, with an author name, and a relevant MeSH term (e.g., search set #013 and #017). Practically speaking, this finding indicates that search results may be more uniform across platforms when searching for a known set of articles using a highly specific, multi-faceted search query. Conversely, simple queries using just one or two keywords or MeSH terms appear more susceptible to significant variations across platforms, underscoring the continued importance of advanced training in literature database searching and consultation with trained information professionals.

However, some platforms appear to be simply broken because they are not able to handle exploding the MeSH hierarchy similarly (e.g., EBSCOhost and Web of Science outliers in search set #008), or they drop records from one month to the next even though the query has not been altered. The lack of discernible causes of significant variance in search result counts over time makes it impossible to adjust for such variance and undermines the trust in using bibliographic databases to inform data-driven decision making.

Our longitudinal study suggested that some differences might be attributed to delays in transferring the MEDLINE file to the vendors, since PubMed updates MEDLINE daily but the other vendors may receive that update and then add that update at later dates. To test this, we ran a COVID-19 search based on a query provided by the NLM in January 2020 and found that there were uneven search result hits for new literature on the COVID-19 pandemic across platforms. Although some of the differences in search result counts might be explained by the previous issues, the main explanation here is likely due to delays in receiving and incorporating the PubMed updates to the vendors. This suggests that if researchers need urgent access to the timely publications, they should be concerned about which version of PubMed they use to collect data.

## Conclusion

Remarkably, these results suggest we may be able to level one of the early critiques of Google Scholar, which was its inability to replicate results from the same search over periods of time [56], on MEDLINE. What followed with research on Google Scholar were several studies recommending against using Google Scholar as the sole database for systematic reviews [35,57,58]. If this criticism is valid for the MEDLINE platforms, our results may strengthen the recommendation by Cochrane [23] that no single MEDLINE platform should be considered a sole source.

The MEDLINE data is licensed to multiple vendors of information services who provide access to the database on their platforms, such as EBSCOhost, ProQuest, Ovid, and Web of Science. Any of these platforms are used by information specialists, health science librarians, medical researchers, and others to conduct research, such as systematic reviews, in the biomedical sciences. Our research examines results based on 29 sets of 145 carefully constructed search queries, plus queries related to the COVID-19 pandemic, across these platforms and indicates that these platforms provide uneven access to the literature, and thus depending on the platform used, the validity of research based on the data gathered from them may be affected. Additional research is required to understand other search related differences among these platforms, including differences among the records that are retrieved, and how they specifically impact research designs like systematic reviews and other biomedical research, and scientific conclusions based on these studies.

